# The use of a centrifuge-free RABiT-II system for high-throughput micronucleus analysis

**DOI:** 10.1101/627596

**Authors:** Mikhail Repin, Sergey Pampou, David J. Brenner, Guy Garty

## Abstract

The cytokinesis-block micronucleus (CBMN) assay is considered as the most suitable biodosimetry method for automation. Previously, we automated this assay on a commercial robotic biotech high-throughput system (RABiT-II) adopting both a traditional and an accelerated micronucleus protocol, both using centrifugation steps for lymphocyte harvesting and washing, after whole blood culturing. Here we describe further development of our accelerated CBMN assay protocol for using on High Throughput/High Content Screening (HTS/HCS) robotic systems without a centrifuge. This opens the way for implementation of the CBMN assay on a wider range of commercial automated HTS/HCS systems and thus increases the potential capacity of dose estimates following a mass-casualty radiological event.

## Introduction

In the case of a mass-casualty radiation event, it is necessary to estimate radiation doses to hundreds of thousands of individuals [1]. On smaller scales this is done manually via one of the IAEA-recommended [2] well-established cytogenetics assays: the dicentric chromosome assay (DCA) or the cytokinesis-block micronucleus (CBMN) assay [3–5]. However, at larger scales, manual analysis of blood samples, even using a large cytogenetic network of labs [6, 7] becomes infeasible and an increase in overall throughput via automation of triage radiation biodosimetry is needed. The CBMN assay is the most suitable biodosimetry method for automation [8, 9].

While custom robotic systems have been built by us [10] and others [11] for radiation biodosimetry, robotic systems face reliability issues unless in continuous use. On the other hand, universal biotech high-throughput/high content screening (HTS/HCS) robotic systems [12] can switch between several programs, offering a high degree of flexibility for using for different high-throughput biological assays. These systems are in routine use for different purposes [13] ensuring their reliability if needed to respond to a radiological event.

Previously, we proposed to use of commercial universal biotech robotic systems for automated preparation of blood samples and automated imaging in ANSI/SLAS-compatible plates [9, 14, 15]. We called this approach RABiT-II (Rapid Automated Biodosimetry Technology – RABiT). Our driving philosophy being that the protocols we develop should be usable on any HTS/HCS system. Seeing that many commercial robotic systems are not fitted with an automated centrifuge, as they are designed for assays that do not require centrifugation (for example, preparation and analysis of attached cell lines). Such systems cannot be exploited for preparation of CBMN assay samples using traditional protocols, which rely heavily on centrifugation [2] for washing of non-adherent cells, thus limiting the number of potential RABiT-II systems. The development of a centrifuge-free automated assay could represent a solution for the use of centrifuge-free robotic systems for preparation of samples for CBMN assay and increase the overall throughout of dose estimate in the case of a large radiological event.

Here we describe the development of the accelerated CBMN assay protocol for sample preparation on a commercial robotic system that does not have a centrifuge (Fig. 1). As a starting point we used the automated accelerated CBMN assay with the reduced time of cell culturing 54h [15], we have previously implemented on a cell::explorer system (PerkinElmer, Waltham MA) which has an integrated, automated centrifuge. In that assay we were able to reduce the time to answer of the conventional CBMN assay by 16 h.

**Figure 1.**
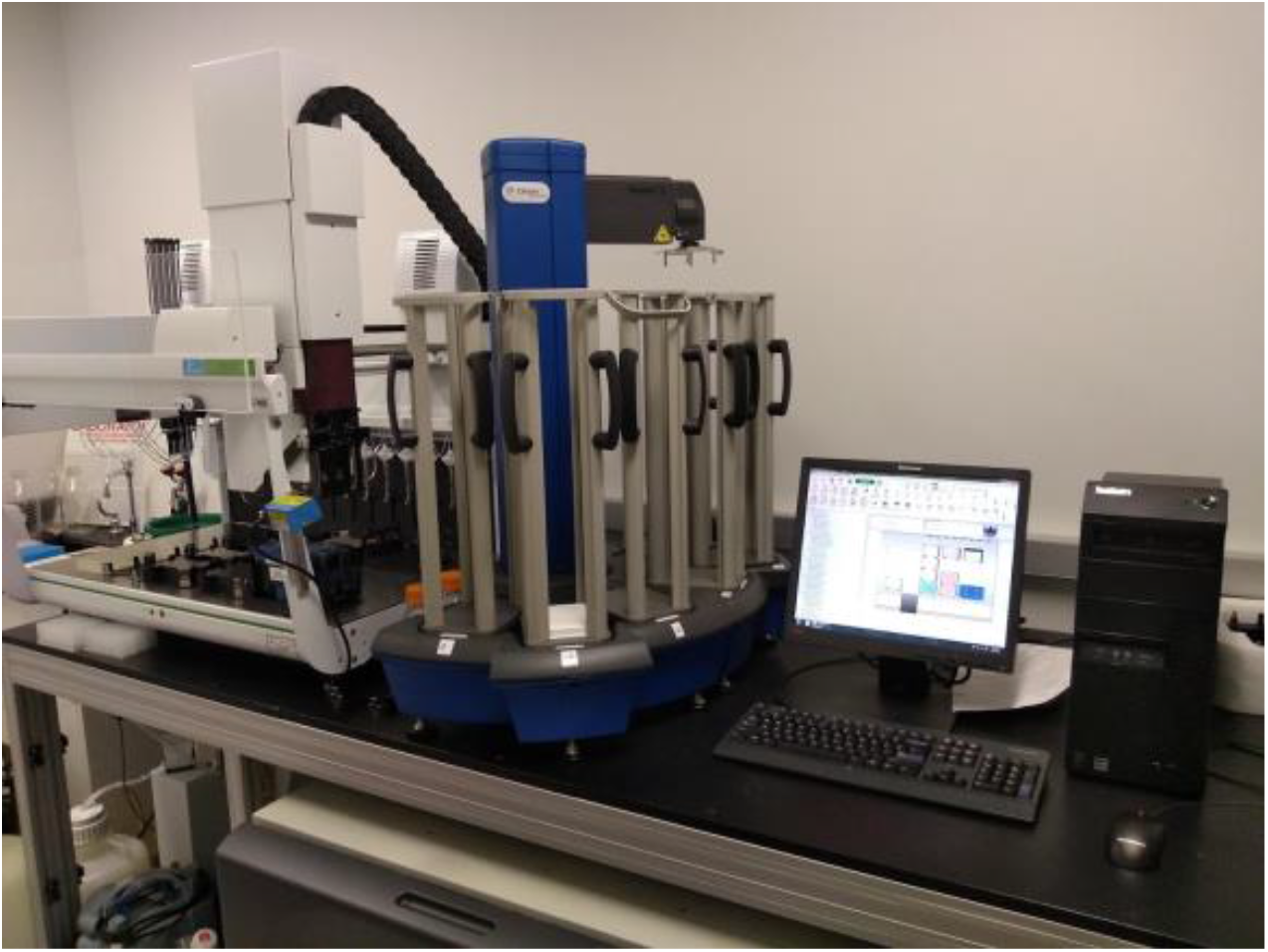
Centrifuge-free RABiT-II system for preparation of samples with automated liquid-handling system JANUS (PerkinElmer) and universal microplate handler Twister3 (Caliper Life Sciences).

## Materials and methods

### Blood sample collection, irradiation and culturing

Blood samples (2 ml) were collected into heparinized vacutainer tubes (BD, Franklin Lakes, NJ) from 4 healthy volunteers following informed consent (IRB protocol #AAAF2671). 24 aliquots of human blood (20 μL) from each donor were pipetted into 1 mL 2D-barcoded tubes (Matrix Storage Tubes; Thermo Fisher Scientific Inc., Waltham, MA), placed into a ANSI/SLAS microplate format compatible 96-tube racks (8 × 12) and covered with the supplied top (Thermo Fisher Scientific). The covered tubes in racks were transported to a Gammacell 40 ^137^Cs irradiator (Atomic Energy of Canada Ltd., Mississauga, Canada). Blood samples were exposed to 0 (control), 1.0, 2.0, 3.0, 4.0 or 5.0 Gy of γ-rays at a dose rate of 0.70 Gy/min. All 96 blood samples (4 sets from each donor) were collected into one 96-tube rack and 200 μL of PB-MAX™ karyotyping media (Thermo Fisher Scientific) was added to each 20 μL blood sample. The rack of tubes containing blood and PB-MAX™ karyotyping media was placed into the incubator at 37°C under 5% CO_2_ atmosphere. After 24 h of incubation, cytochalasin-B (Sigma-Aldrich, St. Louis, MO) was added to the cultures to block cytokinesis of proliferating lymphocytes at a final concentration of 6 μg/mL and cells were cultured for an additional 30 h (total incubation time − 54 h).

### Automated centrifuge-free sample processing after culturing

A centrifuge-free RABiT-II system was used for automated sample processing after culturing: The system consists of two part: the first (Figure 1) has an automated liquid-handling system (JANUS, PerkinElmer) with integrated automated microplate handler (Twister3, Caliper Life Sciences, Waltham, MA) and was used for sample preparation without centrifuge; the second part, an IN Cell Analyzer 2000 automated imager (GE Healthcare, Chicago, IL), was used for sample imaging.

The overall scheme of automated cell harvesting after whole blood culturing is shown in Figure 2. After completion of cell culturing, all the samples from one 96 tubes were divided into two equal parts by transferring to two standard height glass bottom 96 square wells plates (630 μL well volume; Brooks Automation, Chelmsford, MA) preloaded with 300 μL of hypotonic solution (0.075 M potassium chloride). After 1.5 h of sedimentation of cells, the liquid was aspirated leaving ~100 μL, 300 μL of the same hypotonic solution was added and mixed. After another 1.5 h of cell sedimentation, 100 μL of freshly prepared fixative (3:1 methanol:acetic acid) was added from the top. After 10 min, the fixative (3:1 methanol:acetic acid) was exchanged four times using a 200 μL volume, leaving ~100 μL of liquid in each well after each aspiration step. After that, the fixative was exchanged with 200 μL of fixative with increased percentage of methanol (10:1 methanol:acetic acid) and the liquid was fully aspirated. The imaging plates were kept for 30 min for complete evaporation of the residual fixative. For DNA staining, 200 μL of PBS containing 1.5 μg/mL DAPI (4’,6-diamidino-2-phenylindole; Thermo Fisher Scientific) was added for 30 min and exchanged to PBS. All reagents were stored on the deck of the automated liquid-handling system at room temperature.

**Figure 2.**
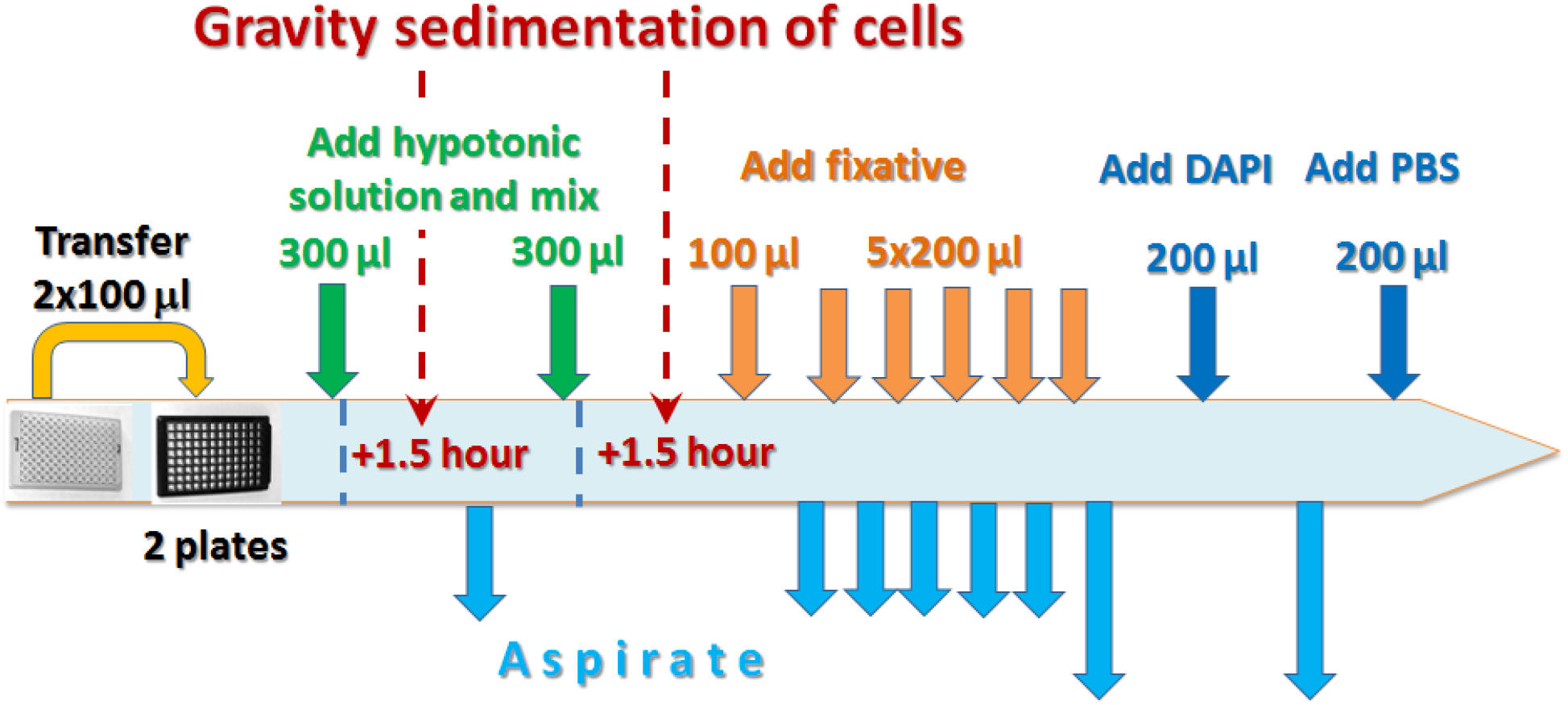
Overall scheme of the centrifuge-free accelerated CBMN assay protocol for samples preparation after whole blood culturing.

### Automated imaging and data analysis

Glass-bottom microplates with fixed and stained samples were scanned on an IN Cell Analyzer 2000. TIFF-images were acquired through a 20x lens with CCD camera (2,048 × 2,048 pixel resolution) using 350/50 nm excitation and 455/58 nm emission at an exposure time of 300 ms with laser autofocusing. Full well image acquisition produced 81 images for each 60 mm^2^ well.

Images were analyzed using the Developer Toolbox 1.9 software (GE Healthcare). Round objects of >0.5 μm granule size (the length of the smallest cross-section through the smallest granule) were identified as micronuclei using the vesicle segmentation module and round objects of >70 μm^2^ were identified as nuclei using the nuclear segmentation module. The data for these identified objects were exported into Microsoft Excel 2010 for further data analysis. Binucleated cells were identified with the following criterion: two nuclei, each with an area larger than 70 μm^2^, similar object characteristics (brightness and area are within 5% and 20%, correspondingly), and with the distance between them less than three times their average radius. A micronucleus was scored if the distance from its center to the midpoint between the centers of two nuclei in the associated binucleated (BN) cell was less than 13 μm, and also that the area of the micronucleus was less than 1/3 of the average area of the two nuclei in the associated BN cell.

## Results

A typical image showing one field of view in a control human blood sample, processed using the centrifuge-free RABiT-II assay with 54 h of cell incubation is shown in Figure 3.

**Figure 3.**
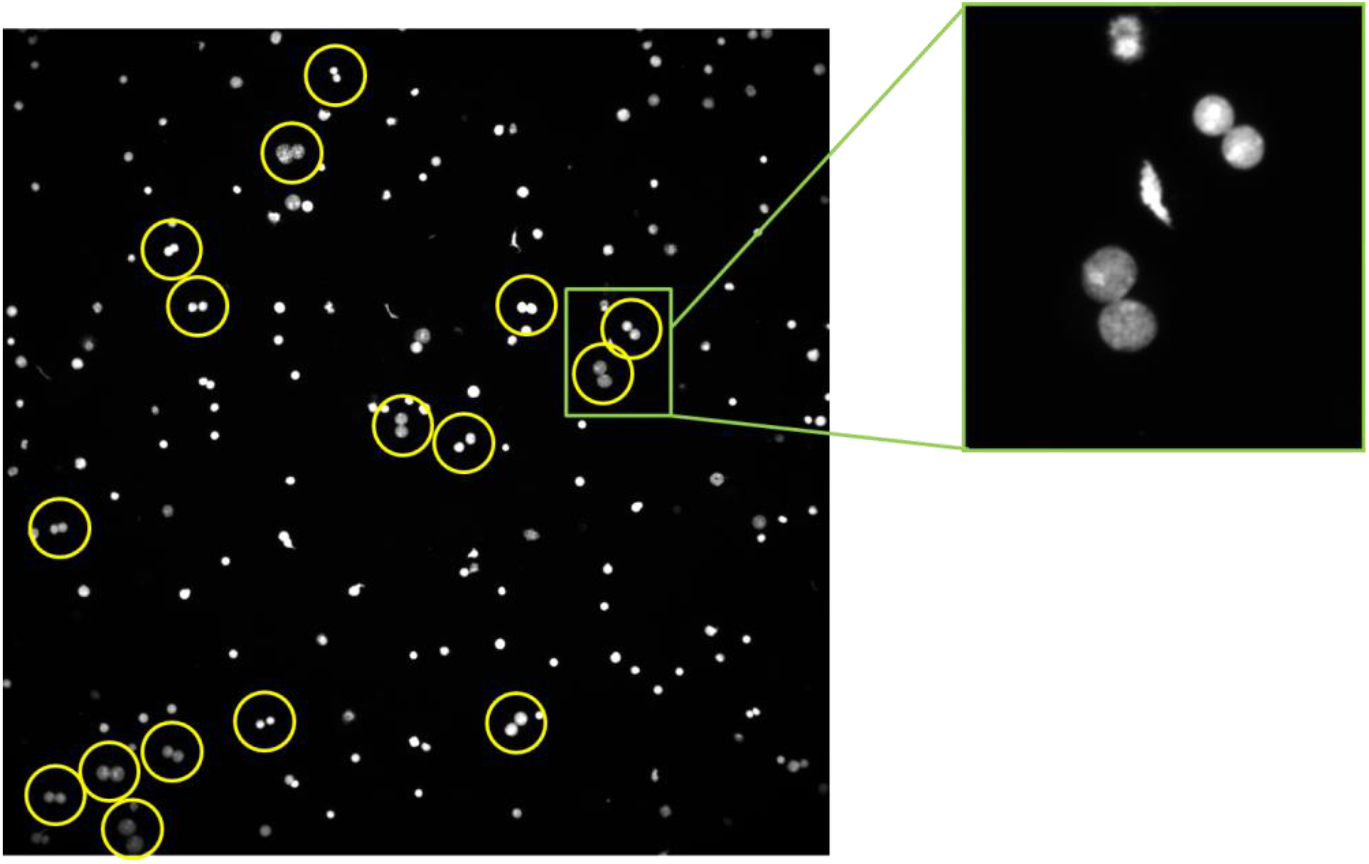
Image of the centrifuge-free RABiT-II accelerated CBMN assay control sample with binucleated cells.

The number of detected BN cells varies with a maximum of 953 and average of 419 in control samples and minimum of 73 BN cells and average of 117 for samples irradiated with the highest dose of 5.0 Gy.

The dose-response curves of micronuclei yields for four healthy donors are shown in Figure 4.

**Figure 4.**
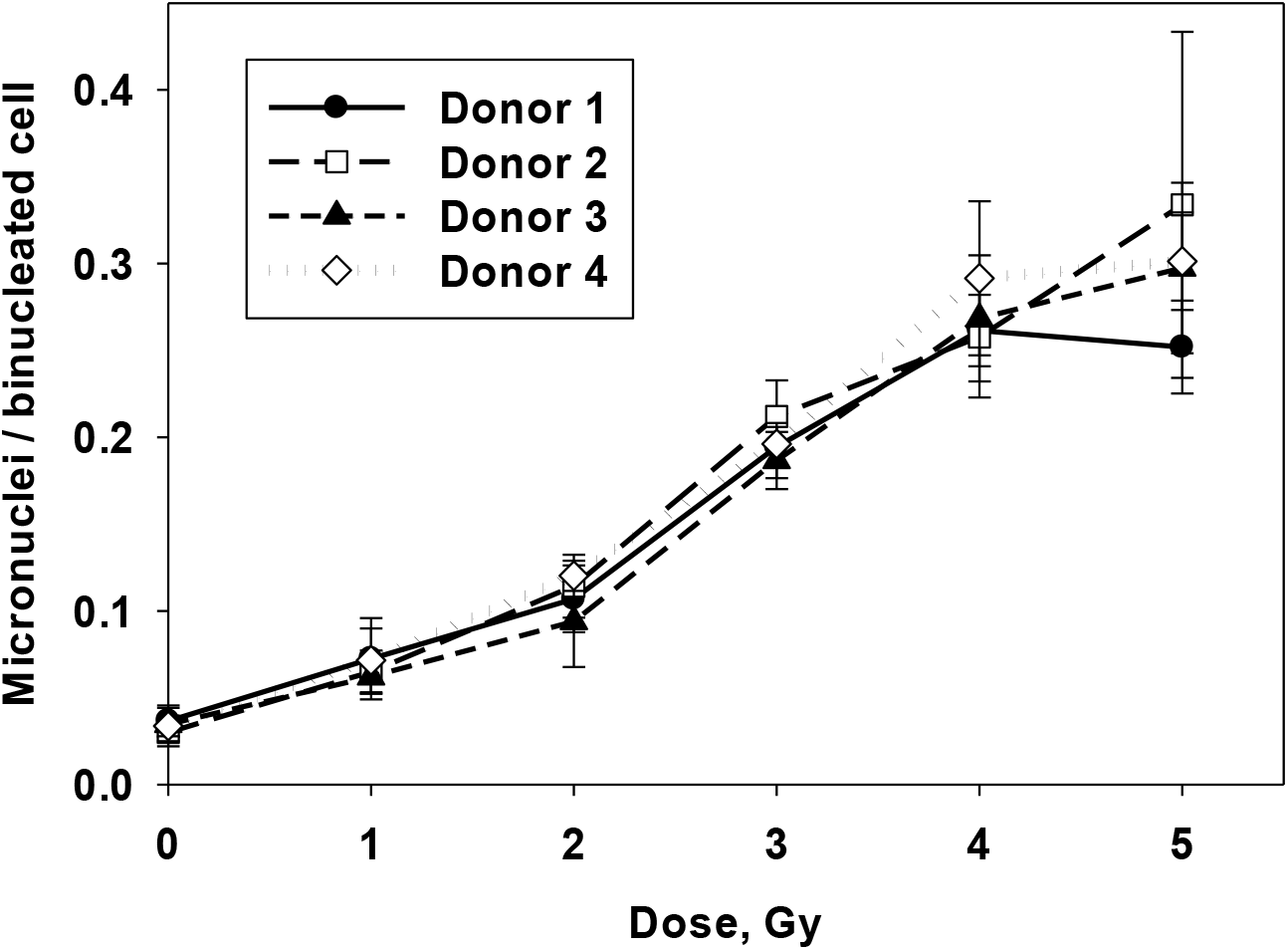
Radiation dose-response curves for four healthy donors obtained by using centrifuge-free RABiT-II accelerated CBMN assay. Error bars correspond to standard deviation of 4 identical samples per donor, processed on the same

## Discussion

Centrifugation is a traditional technique for sample preparation in human cytogenetics and centrifugation-based methods are recommended for preparation of samples in radiation biological dosimetry [2]. In this paper, we show that appropriate samples for the CBMN assay can be obtained by using automated high-throughput HTS/HCS robotic system without a centrifuge. This approach is based on the substitution of multiple centrifugation steps for sample preparation after whole blood culturing to only two steps of gravity sedimentation of cells in standard height imaging microplates and fast multiple washes after adhering of cells to the glass bottom of microplate wells.

The most critical step in the developing of the centrifuge-free protocol was to attach non-adherent lymphocytes to the bottom of wells. For this purpose we used low speed dispensing and aspiration of liquid before the complete attachment of cells following the last aspiration of fixative and its evaporation. We used the sedimentation of cells twice instead of one time before addition of fixative to decrease the amount of debris remaining on the glass bottom of microplate wells before addition of fixative to an acceptable level for further image analysis.

After the second sedimentation of cells, fixative is added to the sample to adhere cells to the glass surface. By changing the standard fixative (3:1 methanol:acetic acid) the purity of samples increases with each step. For the last washing step with fixative solution, the increased concentration of methanol (10:1 methanol:acetic acid) was used to increase the quality of cell preparation for image analysis, particularly for improving the separation of micronuclei from main nuclei in BN cells. Following complete evaporation, DAPI solution is used to stain nuclei content of cells and changed to PBS after staining to increase the contrast of images due to larger debris remained after such procedure as compared with preparation of sample using centrifugation steps.

The quality of CBMN assay samples prepared using the developed centrifuge-free RABiT-II approach (Figure 3) allows to reproduce typical dose-response curves using automated image analysis (Figure 4). This is our first results of centrifuge-free RABiT-II CBMN assay. The developed protocol can be further modified to improve the quality of cells for image analysis. For example, the cell purification step with using magnet-beads can be introduced or with the use of antibody capturing of different primary human lymphocytes as it was demonstrated for CBMN assay on microarray platform [16].

The time of sample preparation after culturing using centrifuge-free robotic system was about 4 h for 96 samples including two gravity sedimentation steps of 1.5 h each. The use of these two sedimentation steps instead of centrifugation steps certainly increases the time of the CBMN assay, but keeps the total assay time within the same 3 days as in the case of accelerated centrifuge-based RABiT-II CBMN assay [15] including cell culturing, image and data analysis. However, the introduction of such centrifuge-free approach and use of the corresponding RABiT-II systems in different laboratories and companies potentially increases the overall throughput of samples preparation by tens of times.

Moreover, our protocols for RABiT-II cell harvesting with the use of centrifuge [9, 15, 17] as well as described in this paper centrifuge-free RABiT-II approach were developed for keeping all reagents and plates at room temperature so that more workstations without automated heaters/coolers could be used for CBMN assay sample preparation. Thus, almost any ANSI/SLAC-format compatible robotic cell handling system meeting the minimum requirements of the centrifuge-free RABiT-II approach can be used for sample preparation with the use of the corresponding RABiT-II systems has the potential to increase the capacity for biodosimetry response during a large-scale radiological/nuclear event.

## Conclusion

The results of this work demonstrate that the accelerated CBMN assay can be automated in a high throughput format by using centrifuge-free commercial HTS/HCS robotic systems designed for running biological assays in standardized multiwell plates. The usage of centrifuge-free biotech robotics as RABIT-II systems in multiple laboratories could considerably increase the total throughput of dose estimations in emergency triage radiation biodosimetry.

